# Modulation of emotion perception and memory via sub-threshold amygdala stimulation in humans

**DOI:** 10.1101/638767

**Authors:** Krzysztof A. Bujarski, Yinchen Song, Sophia I. Kolankiewicz, Gabriella H. Wozniak, Angeline S. Andrews, Sean A. Guillory, Dave W. Roberts, Joshua P. Aronson, Barbara C. Jobst

## Abstract

A common human experience is noticing that emotional life events are more vividly remembered than dull ones. Studies show that the amygdala plays a central role in such emotionally driven enhancement of memory. With this in mind, we investigated the effect of electrical brain stimulation of the left human amygdala on performance on an emotional perception and emotional memory task. We randomly applied sub-threshold 50 Hz stimulation to the left amygdala in 10 patients (5 female and 5 male) with intracranial electrodes during the encoding portion of an emotional valence perception and emotional memory task. We found that amygdala stimulation did not affect reported valence for neutral stimuli (non-stimulated group average valence 5.34, stimulated 5.38, p=0.68) but it did affect positively (non-stimulated group average valence 7.31, stimulated 6.70, p=0.004) and negatively (non-stimulated group average valence 2.79, stimulated 3.55, p=0.0002) valenced stimuli in effect reporting both valence categories as more neutral. Furthermore, we found that stimulation did not significantly disrupt memory for neutral stimuli (68% vs. 61% correctly remembered p=0.48) or positive stimuli (87% vs. 70% correct, trend towards significant difference p=0.09) but did for negative stimuli (83% vs. 67% correct, p=0.03). These results suggest that electrical brain stimulation by our parameters likely reversibly inhibits amygdala function disrupting neural networks responsible for emotional perception and memory. This effect may have clinical implications in treatment of certain neuropsychiatric disorders, such as emotional dysregulation and post-traumatic stress disorder.

**Statement of significance:** The current study builds and expands on extensive prior research into the function of the human amygdala. It provides the first systematic description in humans of a cognitive change brought about by direct electrical stimulation of the amygdala on perception of emotional valence and emotional memory. The results provide further evidence on the importance of the amygdala in human cognition. Likewise, out method utilized to study the function of the amygdala can be extended to study the function of other brain regions in humans, such as the cingulate. While these results are preliminary and need to be duplicated, we aim to further study the effects of amygdala stimulation on emotional processing including possible therapeutic application for diverse group of neuropsychiatric conditions.

## Introduction

Current evidence supports the view that the human amygdala plays a dual role in processing of both emotion and of memory. With regards to emotion, multiple human functional studies clearly show that the amygdala preferentially activates during perception of stimuli with emotional content (Garavan et al. 2001; Salzman and Fusi 2010; Phelps and LeDoux 2005; Hariri et al. 2002; Sergerie, Chochol, and Armony 2008; Ball et al. 2009; Canli et al. 2000). In addition, studies in patients with destructive amygdala lesions show impairment in perception of emotional stimuli, especially negatively valenced stimuli depicting fear (Anderson and Phelps 2001; Phelps et al. 1998; LaBar and Phelps 1998; Adolphs et al. 1997). With regards to memory, current evidence suggests that interactions between the amygdala, the hippocampus, and the entorhinal cortex are the main mechanisms behind emotional memory, i.e. the fact that certain emotional life events are remembered better than non-emotional ones (LaBar and Cabeza 2006; Hamann et al. 1999; Canli et al. 2000; Kilpatrick and Cahill 2003; Kensinger and Corkin 2004; Sergerie, Lepage, and Armony 2006; Richardson, Strange, and Dolan 2004; Phelps 2004). These findings are supported by human lesional studies which show that patients with damaged amygdala have normal long-term declarative memory with only an absence of the expected enhancement of memory by emotion (Cahill et al. 1995; Adolphs et al. 1997; Anderson and Phelps 2001).

An important source of evidence for the function of the human amygdala has come from studies using intracranial EEG and electrical brain stimulation (EBS). Such studies have most commonly been performed in patients undergoing evaluation for epilepsy surgery. Numerous investigators have reported recordings from electrodes inserted directly into the amygdala of event related potentials (ERPs) to emotional stimuli, especially emotional faces (Krolak-Salmon et al. 2004; Pourtois et al. 2010; Sato et al. 2011; Guillory and Bujarski 2014; Halgren et al. 1978; Oya et al. 2002). Furthermore, EBS studies of the amygdala show that both positively (Smith et al. 2006; Meletti et al. 2006; Lanteaume et al. 2007) and negatively (Oya et al. 2002; Naccache et al. 2005; Meletti et al. 2005; Lanteaume et al. 2007) valenced subjective emotional experiences can be induced by amygdala stimulation.

More recently, several investigators have reported effects of amygdala stimulation on perception of emotional intensity and of memory. Bijanki et al. studied a single patient with intracranial electrodes located in the right amygdala and found that stimulation enhanced positive valence during viewing of faces, i.e. the subject perceived faces more positively when stimulation was applied as compared when it was off (Bijanki et al. 2014). Second, Inman et al. reported in fourteen patients that stimulation of the amygdala during perception of neutral stimuli enhanced subsequent delayed memory for the stimuli (Inman et al. 2018). The investigators postulated that stimulation induced interactions between the amygdala, hippocampus, and entorhinal cortex. In both studies, amygdala stimulation produced effects on emotional valence and memory without inducing subjective emotional states in these patients.

Building upon prior work cited above, the objective in this study was to further investigate the effects of amygdala stimulation on performance on an emotional perception and emotional memory task. Our central hypothesis – as gained from previous studies discussed above – was that stimulation of the amygdala would lead to changes in the perceived emotional intensity of stimuli and in turn lead to an effect on delayed recognition, i.e. emotional memory.

## Materials and Methods

### Patients

Consecutive patients at Dartmouth-Hitchcock Medical Center undergoing intracranial EEG evaluation for epilepsy surgery were recruited for to participate in the study. We included both female and male patients who were willing to participate in research, who were able to consent, who had electrodes which localized to the left amygdala on preliminary co-registration, and who had pre-surgical FSIQ greater than 70.

### Electrodes

Electrodes used for the study were either Ad-Tech SEEG electrodes 0.86 mm diameter with 10 contacts per electrode or PMT SEEG electrodes 0.8 mm diameter 8-12 contacts per electrode.

### Task

The task was designed de novo for the purpose of this study (see Figure 1). 144 pictures were selected from the International Affective Picture System (Bradley 2007) (IAPS) using the following criteria: pictures of scenes with people only, IAPS normal control affective valence values between 2 and 8 (1 and 9 were excluded due to extreme nature of some pictures). Due to the fragile nature of our patients, we excluded pictures with the following criteria: stimuli with medical themes (i.e. hospital themes), stimuli with animals, sexuality, and torture. The 144 pictures were divided into three valence categories based on their IAPS ratings: neutral used IAPS value 4.1-6.0, positive 6.1-8.0, and negative 2.0-4.0. The entire picture set was shown and approved by the DHMC IRB. Prior to presentation, each patient was counselled as to the nature of the stimuli, shown sample pictures, and was given a choice to stop the study at any time if they chose to do so. Of the 144 pictures, 48 were randomly chosen to be the encoding set (16 pictures from each emotional valence category) and were used for all patients during Part 1 - Encoding. The remaining 96 pictures (along with the 48 from Part 1) were used for all patients in Part 2 - Delayed Recognition (see Figure 1).

**Figure 1.**
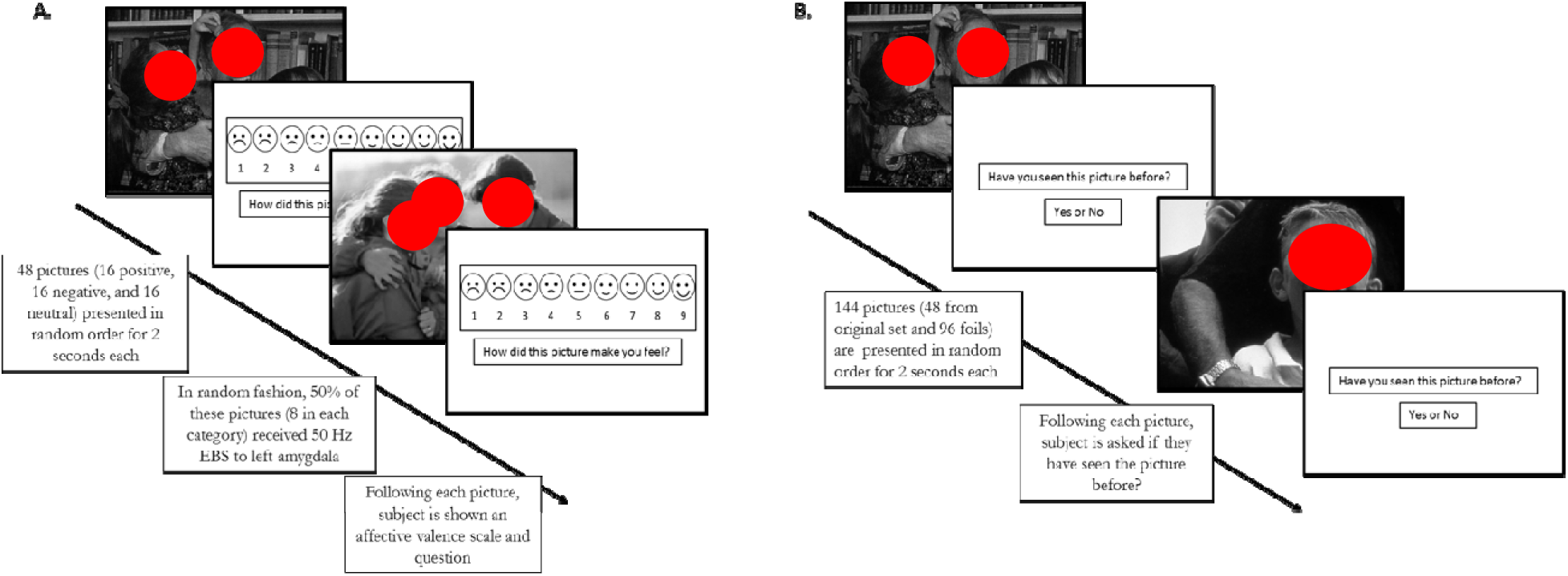
The Affective Valence and Emotional Memory Task

All tasks were administered between June 2015 and April 2018 at Dartmouth-Hitchcock Medical Center in the Epilepsy Monitoring Unit during an approximately 10 day admission for epilepsy surgery evaluation. Following informed consent, a time was chosen when the subject was able to participate in the study. Generally, this occurred when antiepileptic drugs were in the process of being weaned and the patient was awaiting the occurrence of seizures.

### Electrode localization

Coregistration of CT to MRI was done to obtain optimal electrodes for stimulation. Pre-operative whole-brain T1-weighted MRI scans were carried out on a 3-T scanner using a 3D BRAVO sequence with TR = 8.4 ms, TE = 3.4 ms, voxel size = 0.5 * 0.5 * 1.25 mm3. A spine echo T2-weighted coronal MRI scans were also performed to focus on the amygdala and hippocampus areas with TR = 4582 ms, TE = 99 ms, and voxel size of 0.35 × 0.35 × 1.5 mm. All scans were visually inspected by board-certified radiologists for abnormalities, motion and other artifacts. There was no evidence that motion or other artifacts existed in the scans from these ten patients. The left amygdala sub-field segmentation was conducted on the T1-weighted and T2-weighted scans using the Freesurfer tool (Saygin et al. 2017). Post-operative CT head scans were obtained with a voxel size of 0.5*0.5*0.5 mm^3^ to show the locations of depth electrodes. Pre-operative T1-weighted whole-brain MRI scans and post-operative CT head scans were co-registered using SPM12 toolbox in Matlab 2017b. The locations of the depth electrodes were determined by thresholding the registered CT images. Each depth electrode in the registered CT images was represented as a cluster of scattered dots. The coordinate of each depth electrode was calculated by averaging the coordinates of all the dots within each cluster. Individual T1-weighted MRI scans were normalized to Montreal Neurological Institute (MNI) space in SPM12. For visualization purposes, depth electrode coordinates were also transformed to MNI space.

The choice of electrode pair to use for stimulation was made based on the appearance of the co-registration picture. Generally, this included electrodes which were most in the center of the left amygdala. Due to possible distinct roles of the right and left amygdala in cognitive function (i.e. left hemisphere for positive valence, right hemisphere for negative valence) we chose to stimulate the left amygdala for consistency of effect. No further systematic choice of stimulation location was made, i.e. which part of the amygdala.

### Stimulation intensity

The first part of the study included obtaining optimal stimulation parameters for each patient. We defined optimal as a discharge intensity which is safe for clinical applications, did not induce electrographic seizures (afterdischarges) or clinical seizures and did not produce subjective symptoms as gained by patient self-report. A choice of 50 Hz stimulation frequency, 0.3 millisecond pulse duration, and alternating pulse morphology was made for all patients in this study as these parameters are routinely utilized for stimulation mapping at Dartmouth-Hitchcock and are known to be safe for clinical applications. Based on maximum stimulation safety data provided by both electrode manufacturers, we limited maximum stimulation intensity to 5 mA. This stimulation intensity of 5 mA was in the range commonly used for clinical purposes and is well under the 40 microcoulomb per square centimeter stimulation intensity known to possibly cause injury to tissue injury.

### Stimulation paradigm

Initial portion of the experiment involved identifying the optimal stimulation intensity for each patient individually. Starting at 1.0 mA, the chosen electrodes were stimulated in bipolar fashion for 2 seconds observing for electrographic seizures, clinical seizures, and asking the patient to report any subjective symptoms. Duration of 2 seconds was used because this was the duration of picture presentation in the encoding task. If no electrographic seizures, symptoms, or clinical seizures occurred, stimulation intensity was increased stepwise by 1 mA up until either afterdischarges were seen, subjective symptoms were reported by the patient, electrographic seizures occurred (i.e. afterdischarges), or the maximum of 5 mA stimulation intensity was reached. Stimulation intensity was chosen for each patient individually based on 80% of afterdischarge threshold or maximum of 5 mA.

### Task administration

A laptop computer running SuperLab 5 (Cedrus 2015) was connected to a 19.5 inch display monitor which was viewed by the patient at a distance of approximately 3 feet. The subject was given a separate numerical key pad for providing responses. The laptop computer was connected to StimTracker which provided a TTL pulse for triggering the Grass S12X stimulator (FDA approved for clinical use) and separately placing stimulus markers on the EEG. Neuroworks software was used for EEG recording and for electrode selection for stimulation.

For Part 1 – Encoding, the encoding picture set (48 pictures) was presented using SuperLab randomization settings (i.e. by randomization of presented trials in each block). Furthermore, in a random order, 50% of the encoding pictures received electrical stimulation (i.e. by random assignment of “stimulate” or “not stimulate” to each trial). Stimulation of the amygdala was time-locked to the presentation of each picture lasting 2 seconds (same duration as the picture was presented on the screen). Time between presentations of each picture in the encoding task was 10 seconds (i.e. the inter-trial interval).

Part 2 – Delayed Recognition was presented approximately 30 minutes following the initial Part 1 – Encoding. The delay of 30 minutes (as opposed to longer durations such as 24 hours) was chosen between Part 1 and Part 2 for several reasons. First, these patients are apt to experience spontaneous seizures. We aimed to minimize the occurrence of seizures between the two parts of the experiment as seizures can affect cognitive processing and introduce a variable into the results. Second, access to patients with intracranial electrodes in the left amygdala agreeing to participate in research is limited; loss of patients due to inability to perform second portion of the task days later may have limited enrollment. In this part, 144 pictures were presented in random order, 48 from Part 1 - Encoding, and 96 foils. Each picture was presented for 2 seconds and afterwards the patient was asked if they had seen the picture in the initial encoding part 30 minutes earlier.

Patient response data was saved as text files directly from SuperLab and exported to Excel for analysis. For Part 1 – Encoding, each subject’s valence judgments for stimulated and unstimulated pictures were recorded. For Part 2 – Delayed Recognition, each subject’s true positives for stimulated and unstimulated were recorded.

### Data analysis

Data for analysis was exported from SuperLab to Excel for analysis. For Part 1 – Encoding, the results of valence judgments for each picture for all 10 patients was separated into stimulated and unstimulated category. For example, for Patient 1, valence judgments for 24 pictures which received stimulation were given “YES_STIM” category and the rest “NO_STIM”. The same was repeated for Patient 2-10. The exact picture which received stimulation was randomly generated; therefore Patient 2 did not receive stimulation for the same 24 pictures as Patient 1, and so on for Patients 3-10. Average valence per picture for stimulated and unstimulated across all 10 patients was calculated. Pictures were grouped into valence categories as per original picture selection (see picture selection in Methods above). Average valence for stimulated and unstimulated pictures in neutral, positive, and negative categories was calculated and compared using two-tailed T-test and shown in Figure 3B.

**Figure 2.**
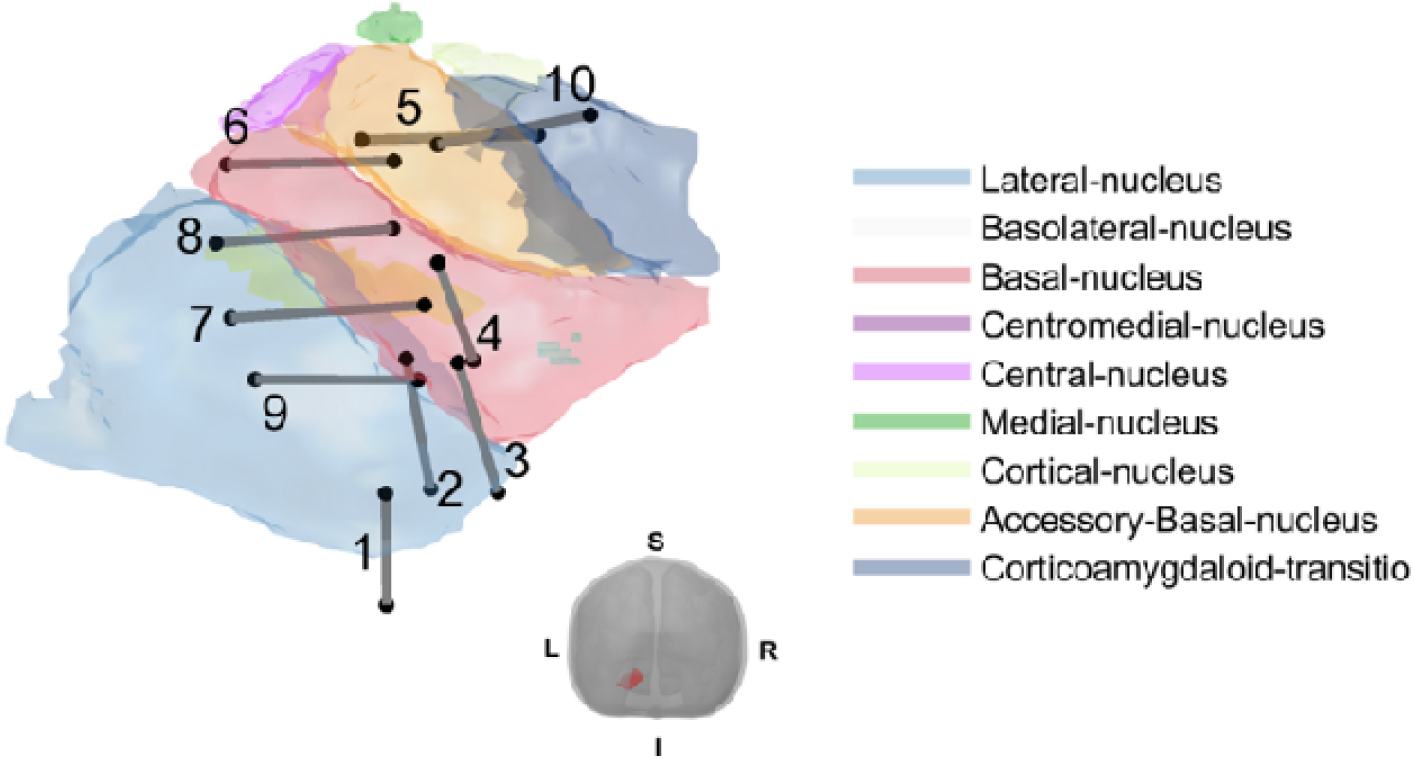
Localization of the stimulating electrode pairs within the left amygdala

**Figure 3.**
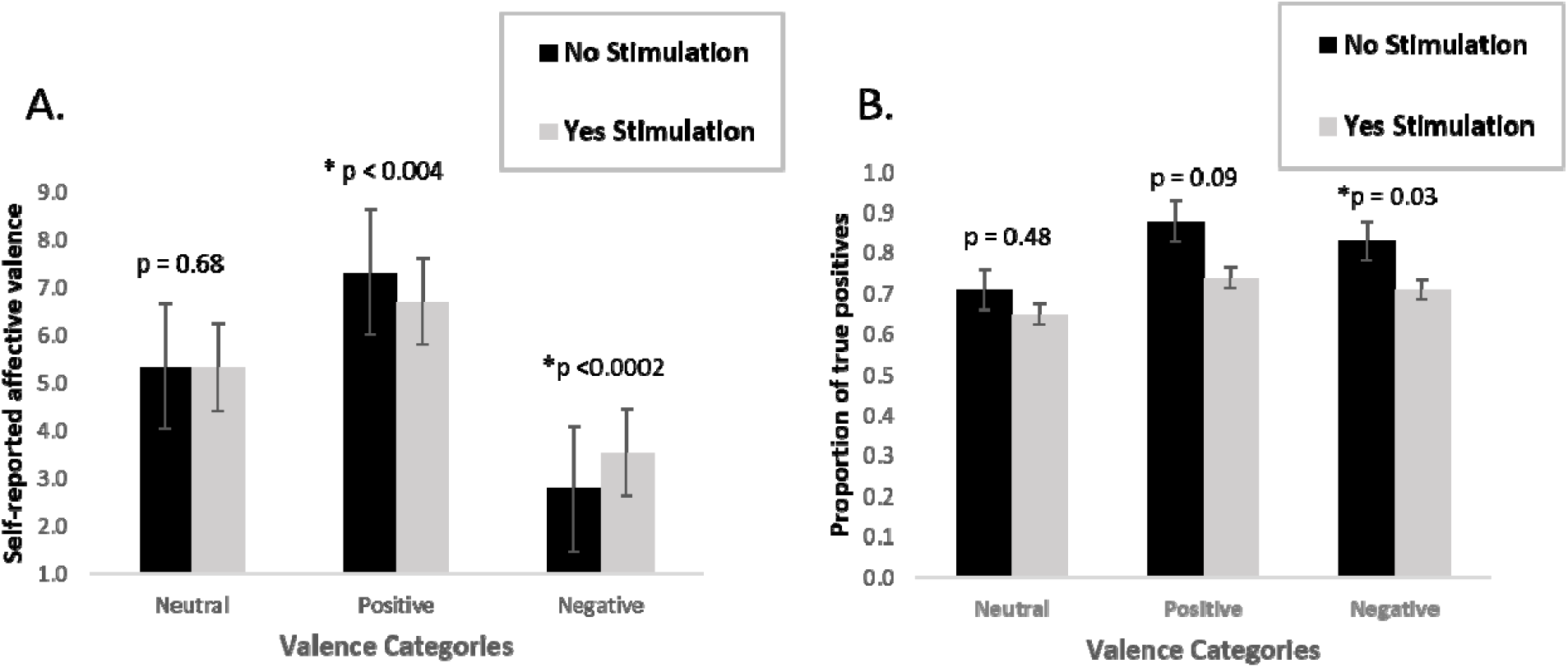
A. Effect of left amygdala EBS on perception of emotional valence for stimulated and unstimulated trials per valence category. B. Effect of left amygdala EBS on 30 minute delayed recognition shown as proportion of true positives for stimulated and unstimulated trials per valence category.

For Part 2 – Delayed Recognition, proportion of true positives was identified for each patient. For example, for Patient 1, of the 144 pictures shown (96 foils and 48 true positives from Part 1 – Encoding) the number of pictures which were correctly identified as having been seen 30 minutes prior was determined and separated per stimulation status (“YES_STIM” vs. “NO_STIM”) and be emotion category (neutral, positive, and negative). Same was done for Patient 2-10. Subsequently, average true positives per emotion category and stimulation status was determined and difference between “YES-STIM” and “NO_STIM” calculated using two-tailed T-test.

## Results

10 consecutive patients who met inclusion criteria are were enrolled into the study. Demographics are shown in Table 1. Patients included men and women of differing epilepsy diagnosis with IQ ranging from 84 to 107 and a variety of seizure onset zones including temporal lobe epilepsy, occipital lobe epilepsy, and frontal lobe epilepsy.

**Table 1.** Demographics.

| SUBJECT | AGE | GENDER | EPILEPSY<br>TYPE | FSIQ | MRI |
| --- | --- | --- | --- | --- | --- |
| 1 | 50 | M | L TLE | 84 | possible L MTS |
| 2 | 41 | F | R TLE | 72 | normal |
| 3 | 22 | F | R TLE | 99 | R MTS |
| 4 | 45 | M | L TLE | 100 | normal |
| 5 | 37 | M | L FLE | 96 | L cingulate dysplasia |
| 6 | 29 | M | L FLE | 102 | normal |
| 7 | 49 | M | L FLE | 107 | left frontal heterotopia |
| 8 | 41 | F | L TLE | 97 | L MTS |
| 9 | 27 | F | L occipital | 100 | periventricular heterotopia |
| 10 | 49 | F | R TLE | 98 | R MTS |

Location of the stimulation electrode pairs are shown in Figure 2. For most patients, both stimulating electrode pairs were within the amygdala, although for several patients one of the electrodes was within the amygdala, while the second was outside of the structure (patients 1, 2, 4). Furthermore, for most patients both electrode pairs were within the basolateral amygdala (patients 1,2,3,4,7,8,9), one patient had contacts in the basolateral and central nuclei (patient 6) and two patients had contacts in the accessory basal nucleus and the corticoamygdaloid transition (patient 5 and 10).

Results of obtaining the optimal amplitude for stimulation are shown in Table 2. In most patients, brief self-limited electrographic seizures or “afterdischarges” were eventually seen with stimulation and the amplitude was reduced to 80% of the “afterdischarge threshold”. Such afterdischarges are common occurrence during functional mapping procedures for clinical applications. The average afterdischarge threshold was 3.4 mA (about 17 microcoulombs per square centimeter of brain tissue) and ranged between 2-5 mA (range of 11.9-29.8 microcoulombs per square centimeter of brain tissue). Two patients self-reported subjective feeling of “nausea” during an afterdischarge (patient 3 and 9). This subjective feeling spontaneously resolved in both patients and was approximately 20 seconds in duration. The EEG in both patients showed a non-evolving rhythmic discharge in theta range which self-terminated about 10 seconds following the end of stimulation. Two patients experienced focal seizures during this part of the experiment (patient 1 and patient 7). Both times, the semiology or clinical manifestations of the seizures mimicked the patient’s own seizure semiology and were obtained ipsilateral to the patient’s own seizures focus. There were no adverse effects of the seizures. Similar to afterdischarges, the occurrence of seizures is common occurrence during stimulation-based functional mapping for clinical applications. The study was successfully completed in both patients who experienced seizures approximately one day later at lower stimulation intensities. Final stimulation parameters did not produce any electrographic seizures or self-reported subjective symptoms by patients during stimulation (see Table 2).

**Table 2.** Obtaining optimal stimulation intensity. Stimulation was performed at 50 Hz for 2 seconds in bipolar fashion using electrode pairs which subsequently were used during Task 1 – Encoding. Stimulation was started at 1 mA and incrementally increased by 1mA to 5 mA. If no afterdischarge threshold was achieved, maximum stimulation of 5 mA was used for the experiment.

| Subject | "Afterdischarge"<br>threshold | Clinical symptoms<br>with<br>"afterdischarge" | Clinical<br>Seizures | Seizure semiology | Final stimulation<br>intensity<br>(milliamps) | Final stimulation intensity<br>(microcoulomb per<br>centimeter square) | Symptoms elicited by<br>final stimulation |
| --- | --- | --- | --- | --- | --- | --- | --- |
| 1 | 4 mA | No | Yes | Fear, scared,<br>agitated 20<br>seconds, end | 3 mA | 17.9 uC/cm<sup>2</sup> | none |
| 2 | 4 mA | No | No | none | 3 mA | 17.9 uC/cm<sup>2</sup> | none |
| 3 | 2 mA | Nausea | No | none | 1 mA | 5.9 uC/cm<sup>2</sup> | none |
| 4 | 4 mA | No | No | none | 3 mA | 17.9 uC/cm<sup>2</sup> | none |
| 5 | 4 mA | No | No | none | 3 mA | 17.9 uC/cm<sup>2</sup> | none |
| 6 | 5 mA | No | No | none | 5 mA | 29.8 uC/cm<sup>2</sup> | none |
| 7 | 2 mA | No | Yes | Unresponsive,<br>automatisms<br>x 45 seconds | 1 mA | 5.9 uC/cm<sup>2</sup> | none |
| 8 | 2 mA | No | No | none | 1 mA | 5.9 uC/cm<sup>2</sup> | none |
| 9 | 2 mA | Nausea | No | none | 1 mA | 5.9 uC/cm<sup>2</sup> | none |
| 10 | 3 mA | No | No | none | 2 mA | 11.9 uC/cm<sup>2</sup> | none |

Figure 3.A. shows the effect of stimulation on perception of emotional valence intensity. Stimulation did not have any significant effect on judgment of emotion valence intensity for pictures in the neutral category (non-stimulated group average valence 5.34, stimulated 5.38, p=0.68). In the positive category, stimulation induced an average decline in reported valence towards neutral (non-stimulated group average valence 7.31, stimulated 6.70, p=0.004). In the negative valence category, stimulation induced an average increase in reported valence towards neutral (non-stimulated group average valence 2.79, stimulated 3.55, p=0.0002).

Figure 3.B. shows effects of stimulation on delayed recognition 30 minutes after presentation of stimuli. For neutral stimuli, 68% were correctly identified when stimulation was not used, 61% if stimulation was used. The percentage correctly identified was lower, although the difference was not significant (p=0.48). For positively valenced stimuli, 87% of non-stimulated pictures were correctly identified after 30 minutes, where as 70% if stimulation was used, the difference showed a trend towards significant difference (p=0.09). For negative stimuli, 83% of non-stimulated pictures were correctly identified after 30 minutes, compared to 67% if stimulation was applied, the difference was significant (p=0.03).

## Discussion

This study had two main findings. First, this study demonstrates that subthreshold (i.e. below afterdischarge threshold and below production of subjective symptoms) 50 Hz unilateral stimulation of the left amygdala reduces perceived emotional intensity for both positively and negatively valenced stimuli with no effect for neutral stimuli. Second, such stimulation during the encoding phase of an emotional memory task as described above has no significant effect on delayed recall of neutrally valenced items, but has significant effect of decreasing memory for emotionally valenced stimuli, especially ones which are negatively valenced. The effects of stimulation on behavior described above are akin to effects of destructive amygdala lesions are likely related to stimulation-induced inhibition of amygdala function and the consequences of such inhibition on neural networks important for perception of emotional valence and emotional memory.

Distinct from the initial conceptualization of the amygdala as a center for representation specific emotions such as for example fear, more recent functional imaging studies show that the amygdala becomes active in vast array of distinct contexts. For instance, the amygdala activates during detection of novelty (Blackford et al. 2010) and during identification of unusual features of stimuli (Herry et al. 2007). Two similar concepts have been put forth to explain the wide ranging functional activation of the amygdala. First, the psychological constructionist perspective as reviewed by Lindquist et al. (Lindquist et al. 2012) argues that the amygdala becomes active when sensory information becomes motivationally salient, irrespective of positive or negative emotional valence. Second, a similar concept puts the amygdala as a peripheral hub of the salience network consisting mainly of the anterior insula and cingulate regions which coordinate the selection of stimuli deserving our attention. (Menon and Uddin 2010; Peters, Dunlop, and Downar 2016). We postulate that the effect of stimulation on perception of both positively and negatively valenced stimuli can best be explained by considering the amygdala as part of the salience network.

Non-human animal studies show that activation of the basolateral nucleus of the amygdala is necessary for normal enhancement of memory by emotion likely by augmenting hippocampal long-term potentiation and affecting hippocampal plasticity (Maren 1999; Barsegyan, McGaugh, and Roozendaal 2014; McIntyre et al. 2003; DaCunha 1999; Akirav and Richter-Levin 2002). Furthermore, non-human animal studies have shown that high-frequency EBS of the basolateral nucleus of the amygdala impairs consolidation of emotional memory and fear learning (Goddard 1964; Kesner and Wilburn 1974; Frey et al. 2001; Gold et al. 1975; Maren 1999; McIntyre et al. 2003; Barsegyan, McGaugh, and Roozendaal 2014). These behavioral effects likely result from tonic depolarization of amygdala neurons during the moment of stimulation and inhibition of their function (Adolphs et al. 1997). Our study suggests that similar to non-human animals, human amygdala EBS using our parameters results in similar cognitive disturbances to those demonstrated in non-human animals and result a behavioral state akin to a functional amygdalectomy. These findings are parallel to hippocampal stimulation studies in human subjects which have shown that stimulation of the hippocampus causes disruption of episodic memory, findings akin to a functional hippocampectomy (Halgren and Wilson 1985; Jacobs et al. 2016).

The effect of electrical stimulation of neural structures on behavior likely relates to the underlying neural organization and the exact parameters of stimulation. For instance, it is well recognized that 50 Hz stimulation of the primary hand area of the human motor cortex results in contralateral hand dystonia, i.e. clinical activation. In contrast, similar 50 Hz stimulation over Broca’s area generally leads to interruption of language function, i.e. clinical inhibition (Havas et al. 2015). This discrepant effect may be related to a difference in the functional organization of the underlying cortex, i.e. relatively parallel organization of pyramidal neurons in the primary motor cortex compared to the non-parallel and overlapping functional units in the language cortex. The structure of the amygdala, i.e. compact grouping of functionally distinct neurons with either corticoid or nuclear histology likely amends well to disruption of function by high-frequency EBS.

Difference in stimulation parameters by Inman et al. and current study likely lead to the apparent differences in outcome, i.e. augmentation vs. disruption of amygdala function. Inman et al. use 0.5 mA pulse lasting 1 second which is composed of eight trains of four pulses at 50 Hz. This type of stimulation, or burst stimulation, has been well demonstrated to induce hippocampal LTP in rat models (Larson, Wong, and Lynch 1986) and replicates typical characteristics of hippocampal physiology, i.e. the theta arousal rhythm (Green and Arduini 1954). Such stimulation has been found to induce hippocampal LTP in non-human animal models if applied to the basolateral amygdala via its strong connections (Bass and Manns 2015). The current study uses non-burst continuous high frequency stimulation, i.e. 2 second 50 Hz train. As discussed above, the physiological effects of such stimulation parameters are uncertain, but are postulated to produce inhibition of the neural structure, or a “reversible lesion”. The best human evidence for the “reversible lesion hypothesis” comes from DBS studies of subcortical structures in Parkinson disease where behavioral effects of high frequency stimulation of > 50 Hz mimics a lesion (Herrington, Cheng, and Eskandar 2016; Dostrovsky and Lozano 2002). Therefore, the distinct effect of amygdala stimulation reported by Inman and the current study are likely based of distinct methods of stimulation used by both. The discrepant results between Inman et al. and the current study suggest that varying the stimulation parameters can lead either to functional augmentation or functional inhibition of amygdala function.

The behavioral effects found in this study are related to unilateral stimulation of the left amygdala. The left amygdala was chosen mostly for consistency of effect as prior studies provide some evidence for specificity in amygdala function (Guillory and Bujarski 2014). Although there is no uniformity in findings, the left amygdala is more likely to activate during emotional tasks as compared to the right (Baas, Aleman, and Kahn 2004). Furthermore, the left amygdala may respond better during positively valenced tasks whereas the right amygdala in negatively valenced ones (Ball et al. 2009). The results of the current study provide evidence that the left amygdala functions during perception of both negative and positive emotional valence.

Many limitations and threats to validity need to be considered. A major limitation of this study are the conclusions which can be reached from electrical brain stimulation. As is common to all stimulation studies, the exact extent of brain tissue affected by stimulation is not known. We have not way to tell if stimulation acted only on the amygdala alone or also on neighboring structures such as the hippocampus, or even distant cerebral networks. Second, we did not assess baseline emotional perception and emotional memory in our cohort. Several studies have found abnormal emotional memory in patients with epilepsy (Munera et al. 2015; Machado Lde, Frank, and Tomaz 2010). Moreover, we do not know the baseline functional status of the amygdala which was stimulated and of the contralateral amygdala to the site of stimulation (i.e. the right amygdala). Therefore, we do not know if the effect of unilateral amygdala stimulation as seen in this cohort with epilepsy would extrapolate to the normal population (i.e. external validity, whether the cause-effect relationship generalizes to the normal population).

A further limitation concerns the design of the delayed recall portion of our task. Much of animal work focused on disruption of long-term memory has used time frames of > 24 hours from encoding to recall. Our recall experiment was presented 30 minutes following encoding. As described in the methods, this was largely done due to the concerns that seizures would occur between the encoding and recall parts of the experiment which may have introduced unacceptable variability in results or dropout. Furthermore, opportunities for testing of patients with intracranial electrodes are limited and fleeting. Because of this design difference, we are not certain how our results can generalized to other work done in this field.

Studies of amygdala stimulation may help to further understand the importance of the amygdala in human cognition. Future studies should use lower stimulation intensities, should aim to understand the differences in effect of continuous and burst stimulation on amygdala function, and should aim to understand the impact of right and left amygdala stimulation. Stimulation to modulate amygdala function in humans may have applications in some neuropsychiatric condition, such as post-traumatic stress disorder or for treatment of emotional disinhibition following traumatic brain injury (Peters, Dunlop, and Downar 2016). At present, we do not have any information to assess safety and validity of such treatments.

## Acknowledgements

No further acknowledgments.

## Supplementary information

We have no supplementary information.

